# Movement of the endoplasmic reticulum is driven by multiple classes of vesicles marked by Rab-GTPases

**DOI:** 10.1101/2024.05.14.592021

**Authors:** Allison Langley, Sarah Abeling-Wang, Erinn Wagner, John Salogiannis

**Affiliations:** Cellular, Molecular, and Biomedical Sciences Graduate Program, University of Vermont; Department of Molecular Physiology and Biophysics, Larner College of Medicine at the University of Vermont, Burlington, VT

## Abstract

Peripheral endoplasmic reticulum (ER) tubules move along microtubules to interact with various organelles through membrane contact sites (MCS). Traditionally, ER moves by either sliding along stable microtubules via molecular motors or attaching to the plus ends of dynamic microtubules through tip attachment complexes (TAC). A recently discovered third process, hitchhiking, involves motile vesicles pulling ER tubules along microtubules. Previous research showed that ER hitchhikes on Rab5- and Rab7-marked endosomes, but it is uncertain if other Rab-vesicles can do the same. In U2OS cells, we screened Rabs for their ability to cotransport with ER tubules and found that ER hitchhikes on post-Golgi vesicles marked by Rab6 (isoforms a and b). Rab6-ER hitchhiking occurs independently of ER-endolysosome contacts and TAC-mediated ER movement. Disrupting either Rab6 or the motility of Rab6-vesicles reduces overall ER movement. Conversely, relocating these vesicles to the cell periphery causes peripheral ER accumulation, indicating that Rab6-vesicle motility is crucial for a subset of ER movements. Proximal post-Golgi vesicles marked by TGN46 are involved in Rab6-ER hitchhiking, while other post-Golgi vesicles (Rabs 8/10/11/13/14) are not essential for ER movement. Our further analysis finds that ER to Golgi vesicles marked by Rab1 are also capable of driving a subset of ER movements. Taken together, our findings suggest that ER hitchhiking on Rab-vesicles is a significant mode of ER movement.

**SIGNIFICANCE STATEMENT:**

- Peripheral endoplasmic reticulum tubules move on microtubules by either attaching to motors (cargo adaptor-mediated), dynamic microtubule-plus ends (tip attachment complexes) or motile vesicles (hitchhiking) but the prevalence of each mode is not clear
- Post-Golgi vesicles marked by Rab6/TGN46 and ER to Golgi vesicles marked by Rab1 drive ER movements
- ER hitchhiking on multiple classes of vesicles (endolysosomal, post-Golgi and ER to Golgi) marked by Rabs plays a prominent role in ER movement

## INTRODUCTION

Organelles, vesicles and macromolecular complexes are driven long distances along microtubules by the molecular motors dynein and kinesin. Precise delivery is critical for many cellular functions including migration, division, and polarized growth (Yarwood *et al*., 2020). The endoplasmic reticulum (ER) extends throughout the entire cell and is arguably the most motile organelle, owing to the fact that peripheral ER tubules (’smooth’ ER) form dynamic membrane contact sites (MCS) with virtually all known classes of organelles (Bonifacino and Neefjes, 2017; Wu *et al*., 2018). Defects in ER-MCS and ER movement lead to a variety of neurological disorders and cancer (Morciano *et al*., 2018; Perkins and Allan, 2021; Wang *et al*., 2023a).

There are multiple modes of ER movement on microtubules. One mechanism is via tip-attachment complexes (TAC), where ER tubules tether to microtubule plus ends and are driven by microtubule polymerization (Waterman-Storer *et al*., 1995; Waterman-Storer and Salmon, 1998). To achieve TAC, the ER resident protein STIM1 interacts with the plus-end binding protein EB1(Grigoriev *et al*., 2008). Other ER-resident proteins such as CLIMP63, Kinectin 1 (KTN1), REEP1 and p180 also interact with microtubules and help shape the ER (Zheng *et al*., 2022; Wang *et al*., 2023b). A second, more prominent mode of ER movement is called sliding, where ER tubules are driven on stable microtubules by molecular motors (Waterman-Storer and Salmon, 1998; Friedman *et al*., 2010). For most organelles, motor-driven movement is primarily achieved via cargo-adaptors. These adaptor proteins localize to organelles to recruit and activate molecular motors along microtubules. For example, TRAK1/2, BicD2, and Hook1/3 adaptors recruit motors to mitochondria, Golgi-derived vesicles, and endosomes, respectively (Fu and Holzbaur, 2014; Reck-Peterson *et al*., 2018; Cross and Dodding, 2019). The identification of ER-specific cargo adaptors has remained elusive. It has been suggested that ER-localized proteins kinectin-1 and protrudin serve as cargo adaptors with kinesin-1 (Kumar *et al*., 1995; Zhang *et al*., 2010; Matsuzaki *et al*., 2011; Petrova *et al*., 2020). However, the significance of the kinesin interaction is unclear given the other prominent roles these proteins play including collaborating with ER-shaping proteins to regulate ER morphology (Chang *et al*., 2013; Raiborg *et al*., 2015; Zheng *et al*., 2022; Wang *et al*., 2023b). Taken together, this suggests that either ER-specific motor-cargo adaptors have yet to be identified or other modes of motor-driven ER movement exist.

An intriguing possibility is that the ER slides on microtubules by “hitchhiking” on motor-driven vesicles. Organelle hitchhiking is a newly described, evolutionarily conserved, mechanism where organelles are propelled by associating with motile vesicles at membrane contact sites (Salogiannis and Reck-Peterson, 2017; Christensen and Reck-Peterson, 2022). Notably, these vesicles are marked by Rab GTPases, master regulators of membrane trafficking, and recruit motors via their own sets of cargo adaptors (Horgan and McCaffrey, 2011; Hutagalung and Novick, 2011). In filamentous fungi, peroxisomes, lipid droplets, and ER hitchhike on Rab5-marked early endosomes (Guimaraes *et al*., 2015; Salogiannis *et al*., 2016, 2021). In mammalian cells, the ER hitchhikes on lysosomes, as well as on Rab5- and Rab7-marked endosomes (Friedman *et al*., 2013; Guo *et al*., 2018; Lu *et al*., 2020; Spits *et al*., 2021; Jang *et al*., 2022). Re-localizing endosomes either by artificial recruitment of motors or by nutrient starvation drastically reorganizes ER morphology, suggesting that ER dynamics are coupled to endosome movement (Lu *et al*., 2020; Spits *et al*., 2021; Jang *et al*., 2022). Taken together, Rab-marked endosome movement is required for a subset of ER movements, but it is unclear if other Rab-vesicles participate in hitchhiking. In support of this possibility, loss-of-function of multiple Rabs (1/5/7/10/18) can lead to defects in ER morphology (Audhya *et al*., 2007; English and Voeltz, 2013a; Gerondopoulos *et al*., 2014; Mateus *et al*., 2018; Rollins and Blankenship, 2023).

We hypothesized that other Rab-vesicles may participate in ER hitchhiking. In the present study, we find that post-Golgi vesicles marked by Rab6 are necessary and sufficient for normal movement of ER. Rab6-vesicles cotransport with ER tubules on stable microtubules independently of ER hitchhiking on Rab7-marked endolysosomes. Most of these events are in the anterograde (kinesin-driven) direction towards the cell periphery. Surprisingly, none of the other five post-Golgi Rab-vesicles participated in ER hitchhiking. Instead, Rab6-ER hitchhiking events are enriched for TGN46, a trans-Golgi marker (Luzio *et al*., 1990). We also find that ER hitchhikes on Rab1 (ER to Golgi) vesicles independently of Rab6. Overall, our data suggests that ER hitchhiking on Rab-vesicles is a prominent mode of microtubule-based movement of the ER.

## RESULTS

### ER cotransports with Rab6-marked vesicles independently of late endosomes

Previous work established that a subset of ER tubules moves by cotransporting with endolysosomal vesicles marked by Rab5 and Rab7 (Friedman *et al*., 2013; Guo *et al*., 2018; Lu *et al*., 2020; Spits *et al*., 2021; Jang *et al*., 2022). We hypothesized that Rabs representing other vesicular pathways are also capable of cotransporting with the tips of ER tubules. We initially tested Rab6, a prominent regulator of Golgi-derived trafficking and the most abundantly expressed Rab in the Golgi (Liu and Storrie, 2015). U2OS cells expressing either mCherry-KDEL or mCherry-Sec61β (ER markers) and eGFP-Rab6a were subjected to two-color live spinning-disk confocal imaging (Figure 1A and Supplementary Figure 1A and Supplemental Movie S1). We find that a subset of ER tubules in the cell periphery cotransport with Rab6a-vesicles (Figure 1B and Supplementary Figure 1B). Similar results were obtained for isoform Rab6b (Figure 1B). Rab6a colocalizes with the vast majority of Rab6b, confirming they occupy overlapping vesicle populations (Supplementary Figure 1C). Interestingly, although a subset (∼25%) of ER tubules led by Rab6 (herein referring to Rab6a or Rab6b) were transient and ended with a quickly retracted ER tubule, most ended with either a fusion event across a polygonal ER junction or running through a polygon (Figure 1A and 1C). Furthermore, the interactions between Rab6 and ER were short-lived, and we often detected Rab6-vesicles alone (i.e. without ER) before and after cotransporting with the ER. We explored the possibility that other organelles could hitchhike on Rab6-vesicles by testing mitochondrial cotransport with Rab6-vesicles. We find that mitochondria move independently of Rab6-vesicles, affirming Rab6-ER cotransport is specific (Figure 1D and 1E).

**Figure 1.**
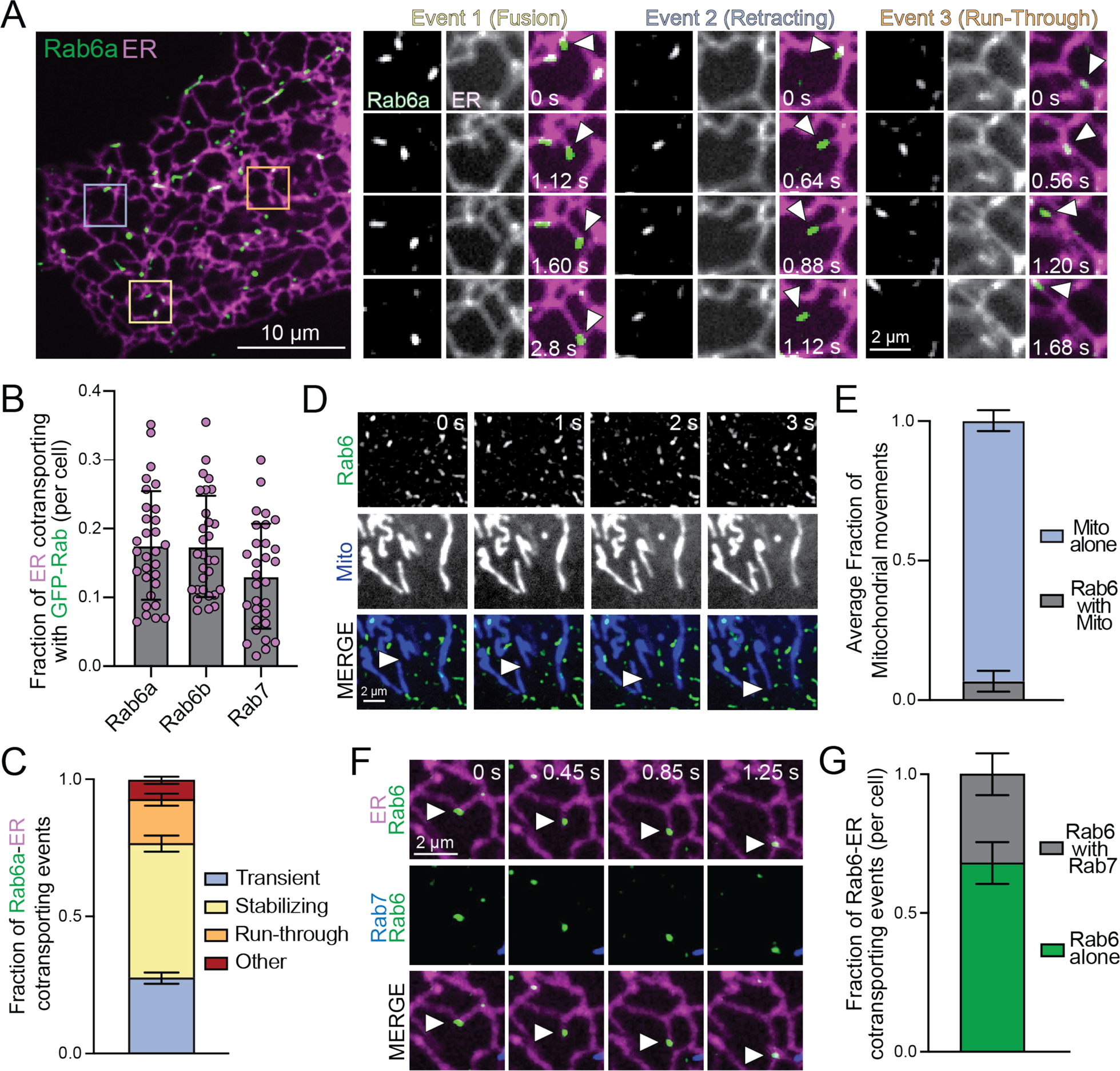
ER tubules cotransport with Rab6-vesicles independently of Rab7-marked late endosomes. (A) U2OS cells expressing GFP-Rab6a and mCherry-KDEL (ER) (left panel) and time-lapse stills (right panels). White arrowheads indicate cotransport events (fusion, retracting, and run-through). Also see Supplemental Movie S1. (B) Cotransport frequency between vesicles and ER tubules in U2OS cells expressing mCherry-KDEL (ER) and either eGFP-Rab6a, eGFP-Rab6b, or eGFP-Rab7. (n=30 cells). Data is mean ± SD. (C) Fraction of different types of ER-Rab6a cotransport events in U2OS cells (n=9 cells). Data is mean ± SEM. (D) Time-lapse stills from U2OS cells expressing Halo-Rab6b (JF549 dye) and dyed with Mito-Tracker Deep Red. White arrowheads indicate mitochondrial movement in the absence of Rab6. (E) Fraction of mitochondrial movements occurring in the presence and absence of Rab6b-vesicles (N=3 technical replicates from 28 analyzed cells). Data is mean ± SEM. (F) Time-lapse stills from U2OS cells expressing mCherry-KDEL (ER), GFP-Rab7, and Halo-Rab6b (JF646). White arrowheads indicate Rab6-ER cotransport without Rab7. Also see Supplemental Movie S2. (G) Fraction of ER-Rab6b cotransport events in the absence and presence of Rab7. (N=5 technical replicates from 32 analyzed cells). Data is mean ± SEM.

Previous work established that Rab7-marked late endosomes (LEs) and lysosomes cotransport with the ER (Guo *et al*., 2018; Lu *et al*., 2020; Spits *et al*., 2021). We find that Rab6-vesicles and Rab7-LEs cotransport with the ER to a similar extent (Figure 1B). We set out to determine if Rab6-ER and Rab7-ER cotransport events are mutually exclusive. Given their distinct roles in trafficking, the majority of Rab6 and Rab7 do not occupy the same vesicles and represent two separate pools (Hutagalung and Novick, 2011; Deffieu *et al*., 2021). We confirmed that Rab6 does not colocalize with Rab7 (Supplementary Figure 1D). In addition, to rule out the possibility that a small subset of motile Rab6/7-occupied vesicles were enriched at ER tubules, we performed three-color live imaging in cells expressing GFP-Rab7, mCherry-KDEL and Halo-Rab6. We found most ER hitchhiking vesicles containing Rab6 were absent for Rab7 (Figure 1F and 1G and Supplemental Movie S2). Taken together, these data suggest that ER tubules can cotransport with Rab6-vesicles independently of Rab7-marked LEs.

### ER tubules hitchhike with Rab6 on stable microtubules and are TAC-independent

The canonical mechanisms of peripheral ER movement are called sliding and tip-attachment complex (TAC). In sliding, ER moves along stable microtubules at speeds consistent with the microtubule-based motors dynein and kinesin in cells (Lee and Chen, 1988; Friedman *et al*., 2010). In TAC, ER speeds are slower due to ER tethering to polymerizing and depolymerizing microtubules (Waterman-Storer *et al*., 1995; Grigoriev *et al*., 2008; Friedman *et al*., 2010). TAC also accounts for the slowest ∼10% of total ER movements (Grigoriev *et al*., 2008; Friedman *et al*., 2010). To determine if Rab6-ER cotransport occurs via TAC or a motor-based mechanism, the speeds of individual ER tubules were quantified with and without an associated Rab6-vesicle. We find that all (49/49) Rab6-ER cotransport events had speeds greater than >800 nm/sec and were absent from the slowest ∼8% of total ER movements (Figure 2A). This is consistent with a TAC-independent mechanism of movement (Waterman-Storer *et al*., 1995; Friedman *et al*., 2010). We also find a slight increase in the speed of Rab6 during a cotransporting event (compared to before and after the event) suggesting no drag on the vesicles (Figure 2B). This contrasts with lysosomes which were previously shown to slow down during ER hitchhiking events (Lu *et al*., 2020). Finally, we fluorescently labeled microtubules (mScarlet-alpha tubulin) in cells expressing mEmerald-sec61β and Halo-Rab6 and find ∼90% of Rab6-ER cotransport events occur on preexisting microtubules (Figure 2C and 2D).

**Figure 2.**
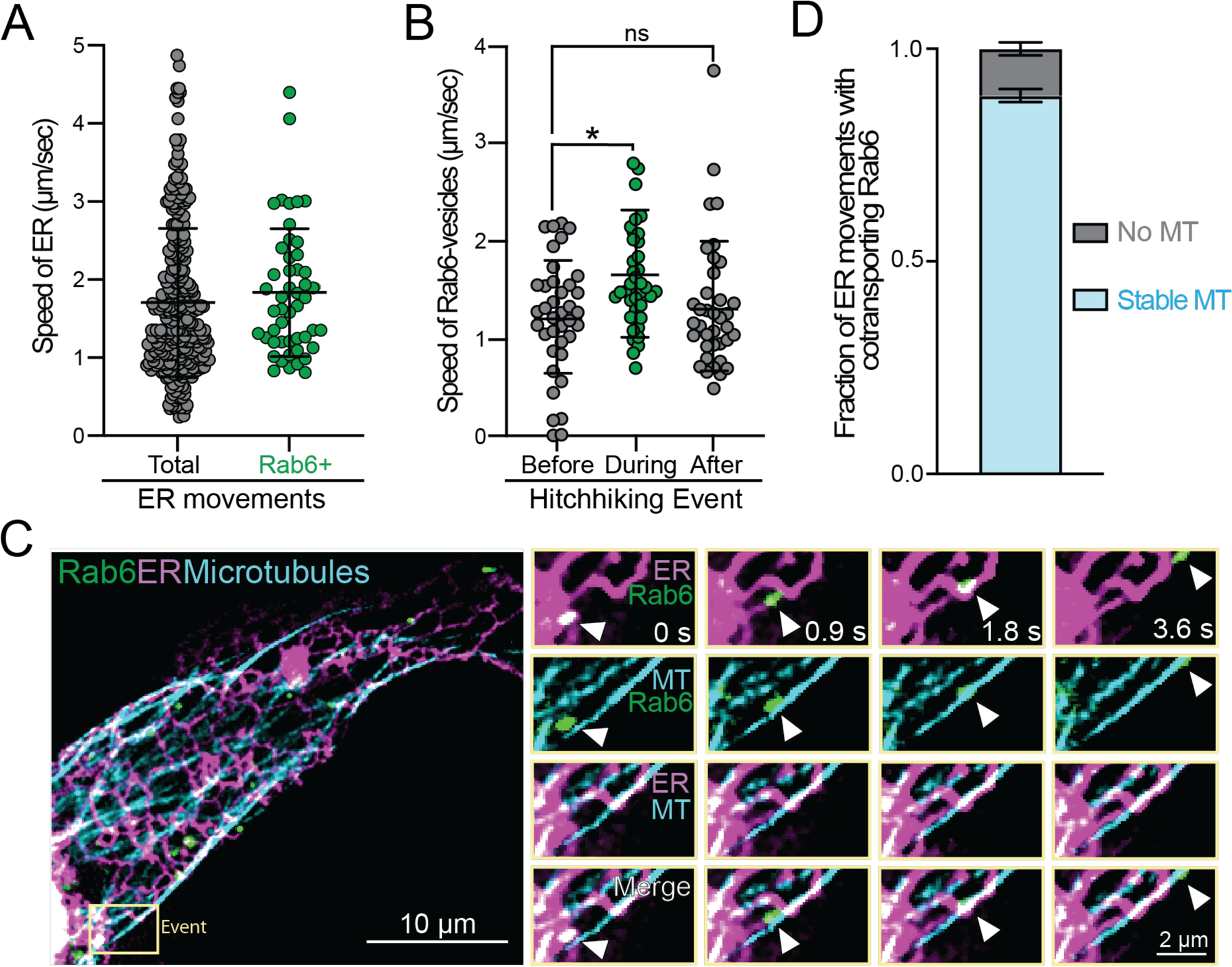
ER-Rab6 hitchhiking occurs on stable microtubules and is TAC-independent. (A) Speed of ER tubule movements with (n=49 events) or without (n=287 events) a cotransporting Rab6-vesicle in U2OS cells expressing GFP-Rab6a and mCherry-KDEL (ER). Data represents mean ± SD. (B) Speed of Rab6a-vesicles before (n=38 events), during (n=39 events), and after (n=37 events) cotransport with ER tubules (Brown-Forsythe and Welch ANOVA, *p<0.05.) Data represents mean ± SD. (C) U2OS expressing mEmerald-Sec61β (ER), mScarlet-tubulin (MTs), and Halo-Rab6b (JF646) (left panel) and time-lapse stills (right panels). White arrowheads indicate ER-Rab6 cotransport events on stable microtubules. (D) Fraction of Rab6-ER cotransport events occurring on stable microtubules (N=3 technical replicates from 20 analyzed cells). Data represents mean ± SEM.

### Rab6-vesicle motility is necessary and sufficient for ER movement

Based on our data thus far, we hypothesize that Rab6-vesicle movements along stable microtubules are coupled to the ER and are necessary for ER movement. To test this initially, we quantified ER movement in cells expressing either GFP-Rab6 WT, constitutively active (CA) GTP-locked (Q72L), or inactive GDP-locked (T27N) dominant-negative (DN) for both Rab6a and Rab6b isoforms. Rab6-WT and Rab6-GTP-locked expressing cells contain punctate and motile GFP-marked vesicles, while Rab6-DN expressing cells have diffuse GFP signal with no discernible vesicles (White *et al*., 1999). DN-expressing cells have reduced ER movements compared to WT and GTP-locked expressing cells (Figure 3A and 3B and Supplementary Figure 2A). Importantly, Rab6-WT had similar ER movement compared to a cell expressing cytosolic GFP, ruling out the possibility of an overexpression phenotype artificially suppressed in the DN conditions (Supplementary Figure 2B). There were also no differences in protein expression levels as confirmed by Western blot (Supplementary Figure 2C and 2D). As an alternative approach, CRISPR/Cas9 was used to generate Rab6a and Rab6b double knockouts (6a/b DKO) in U2OS cells (Figure 3C). ER movement in the periphery was significantly reduced in Rab6a/b DKO cells compared to control cells (Figure 3D).

**Figure 3:**
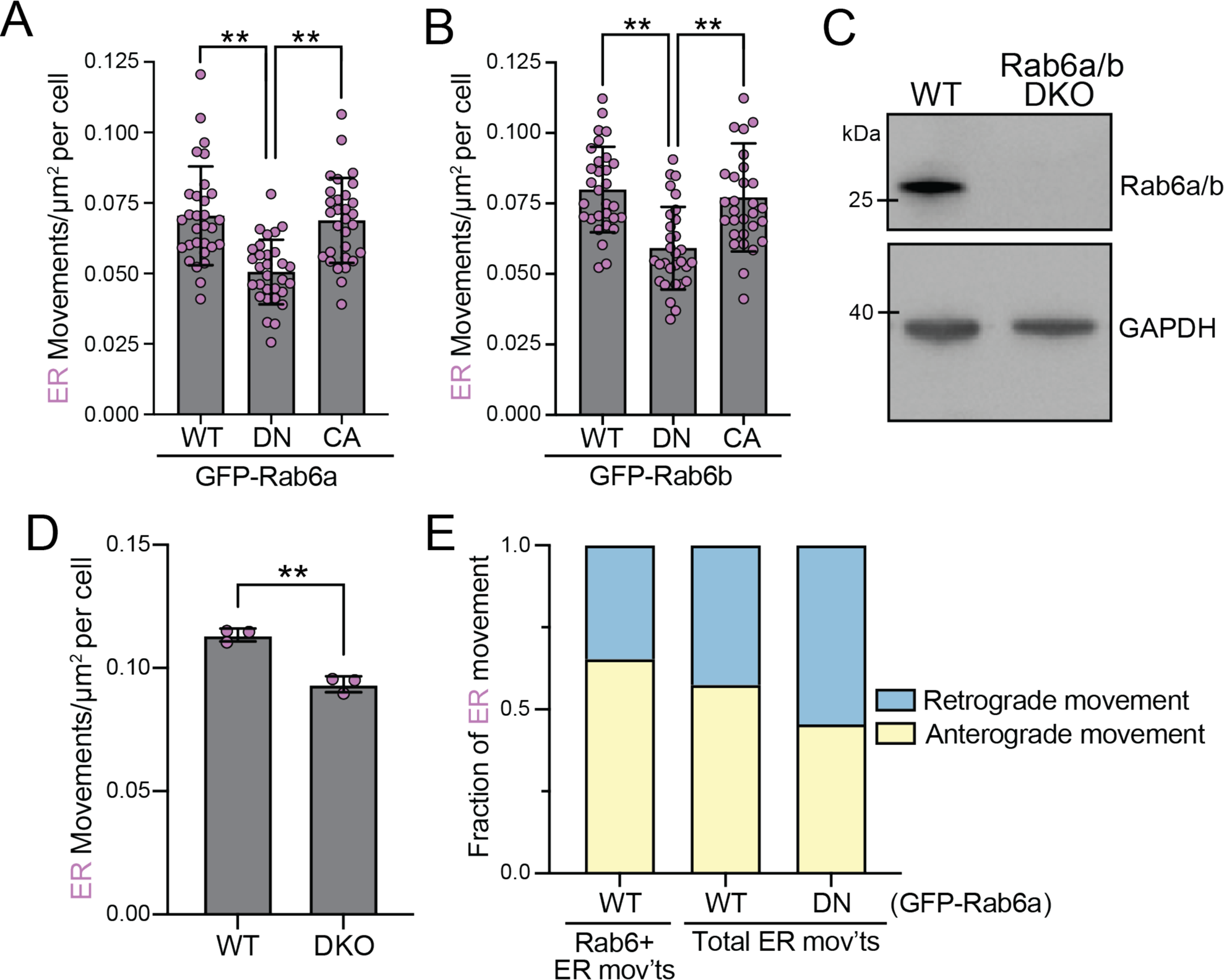
Loss of Rab6 perturbs ER movement. (A-B) ER movement in U2OS cells expressing mCherry-KDEL (ER) with either WT, dominant-negative (DN; T27N; GDP-locked), or constitutively active (CA; Q72L; GTP-locked) eGFP-Rab6a (A) and eGFP-Rab6b (B) (n=30 cells; Brown-Forsythe and Welch ANOVA, **p<0.01). (C) Western blot of GFP-Rab6a/b isoforms from wild-type and double knockout (DKO) CRISPR U2OS cell lines. (D) ER movement in Rab6a/b DKO U2OS cells (N=3 technical replicates from 30 analyzed cells per condition, Unpaired Welch’s t-test, **p<0.01). Data represents mean ± SD. (E) Fraction of ER movements in the anterograde and retrograde direction with and without a cotransporting Rab6-vesicle in U2OS cells expressing WT or DN eGFP-Rab6a (n=287 events for total movements in WT expressing cells, n=49 events for Rab6+ ER movements, n=242 events for total movements in DN-expressing cells; analyzed from same raw data as Figure 2A).

Rab6-vesicles move robustly towards both plus and minus ends in cells, owing to the prominent interactions with kinesins and dynein, respectively (White *et al*., 1999; Grigoriev *et al*., 2007). We set out to determine whether Rab6-ER cotransport events reflected this bidirectional motility or were biased only in one direction. We find ∼65% Rab6-ER cotransport events move in the anterograde direction (towards the cell periphery) (Figure 3E). In addition, total peripheral ER tubules move with a slight anterograde bias, and this bias is suppressed in cells expressing a Rab6-DN mutant (Figure 3E). This suggests that most Rab6-dependent ER movements are driven by plus-end directed kinesins.

The previous loss-of-function experiments were under transient transfection (∼24 hour) or knockout conditions. To test the hypothesis that Rab6-vesicles were required for a subset of ER movements in a more acute manner, we adapted a rapalog-inducible heterodimerization approach to halt vesicle motility while keeping the GTP-binding cycle intact (Figure 4A) (Schlager *et al*., 2014). Two tandem copies of FK506-binding protein (FKBP) were fused to Halo-tagged Rab6 (2xFKBP-Halo-Rab6), and an FKBP-rapamycin-binding (FRB) domain was fused to an mEmerald-tagged truncated kinesin-1 (K560) harboring a T92N rigor mutation (FRB-mEmerald-K560^T92N^), rendering it constitutively anchored to microtubules (Figure 4A). Using this strategy, the movement of Rab6-vesicles was rapidly reduced (< 30 mins) upon addition of rapalog (A/C heterodimerizer) but not with an ethanol control (Figure 4B). We quantified the movement of ER in the same cell before and 30 minutes after treatment. ER movement is significantly reduced after rapalog treatment, but unchanged with ethanol (Figure 4C and 4D). Furthermore, rapalog treatment in cells lacking expression of FRB-mEmerald-K560^T92N^ did not affect ER movement (Supplemental Figure 3A). Taken together, this suggests ER movement is acutely coupled to Rab6-vesicles.

**Figure 4.**
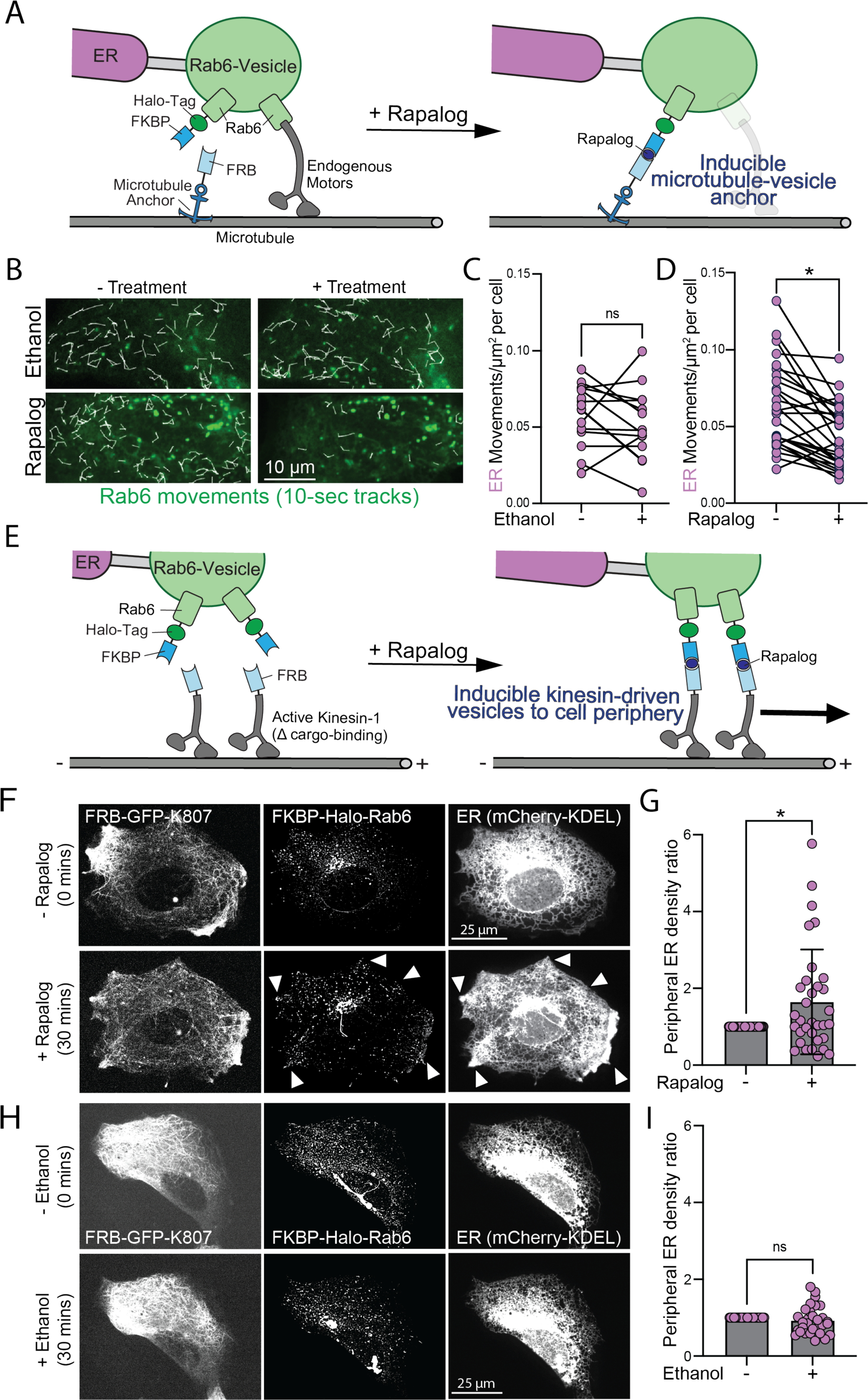
Rapalog-induced approaches demonstrate that Rab6-vesicle movements are necessary and sufficient for ER movement. (A) Schematic of rapalog-induced heterodimerization of FRB-mEmerald-K560^T92N^ with 2xFKBP-Halo-Rab6b for acute stoppage of Rab6-vesicles. (B) Micrograph of TrackMate-generated Rab6b-vesicle tracks in U2OS cells before and 30 minutes after treatment with 5 µM rapalog (microtubule-anchor induction) or ethanol (control). (C-D) ER movements in U2OS cells before and 30 minutes after treatment with ethanol (C) (n=16 cells; Wilcoxon paired test, ns) or rapalog (D) (n=27 cells; Wilcoxon paired test, *p<0.05). (E) Schematic of rapalog-induced heterodimerization of KIF5B^K807^-Emerald-FRB with 2xFKBP-Halo-Rab6b to drive cell periphery directed movement of Rab6-vesicles. (F and H) Micrographs of U2OS cells expressing KIF5B^K807^-mEmerald-FRB, 2xFKBP-Halo-Rab6b and mCherry-KDEL (ER) before and after treatment with rapalog (F) or ethanol (H). White arrowheads indicate areas of Rab6-vesicle and ER accumulation. Also see Supplemental Movie S3. (G and I) Quantification of F and H as a ratio of peripheral to perinuclear ER intensity in the same cell before and after treatment with rapalog (G) (n=33 cells, Wilcoxon paired test, *p<0.05) or ethanol (I) (n=30 cells, Wilcoxon paired test, ns). All data represents mean ± SD. ns=not significant

Next, we determined if Rab6 movement is sufficient to drive ER movement and distribution. To test this, we adapted a rapalog-inducible strategy to drive Rab6-vesicles towards the cell periphery (Kapitein *et al*., 2010; Schlager *et al*., 2014). The FKBP-Halo-Rab6 construct was expressed in U2OS cells alongside an active kinesin-1 lacking a cargo binding domain (KIF5B^K807^) fused to mEmerald-FRB (Figure 4E). With the addition of rapalog, Rab6-vesicles were rapidly (< 30 mins) re-localized towards the cell periphery. Vesicles only localized to a subset of regions along the cell periphery (Figure 4F; white arrows). These enriched regions are consistent with previous reports demonstrating that Rab6-vesicles travel on subsets of microtubules towards secretion hotspots (Fourriere *et al*., 2019). Remarkably, we observed a drastic accumulation of ER at these Rab6 enriched regions (Figure 4F white arrows, and Supplementary Figure 3B and Supplemental Movie S3). We also quantified this as the ratio of peripheral to perinuclear ER intensity in the same cell before and 30-60 minutes after treatment and find that this ratio significantly increased after rapalog treatment (Figure 4G). No distribution changes were observed with ethanol treatment (Figure 4H and 4I).

### Proximal, but not distal, post-Golgi vesicles participate in ER hitchhiking

Motile vesicles marked by Rab6 are Golgi-derived (Grigoriev *et al*., 2007). Specifically, Rab6 regulates cis-Golgi to ER trafficking, intra-Golgi transport, post-Golgi secretory vesicles targeted to the plasma membrane, and trafficking routes destined for and recycled back from endosomes (White *et al*., 1999; Mallard *et al*., 2002; Grigoriev *et al*., 2007; Dickson *et al*., 2020; G. Dornan and C. Simpson, 2023). We set out to determine the specific source of the Rab6-vesicles participating in ER hitchhiking.

Initially, we screened Rab proteins known to be involved in post-Golgi vesicular trafficking (Galea and Simpson, 2015) to determine if they functionally overlap with Rab6 in ER movement. Compared to distal post-Golgi Rabs 8,10 and 11, Rab6 is the only GFP-Rab enriched at the leading tip of ER tubules (Figure 5A). Furthermore, although expression of Rab6-DN disrupts ER movement (Figure 3A and 3B), DN constructs of other post-Golgi Rabs (8/10/11/13/14) did not affect ER movement (Figure 5B and Supplementary Figure 4A). Importantly, total ER movement is similar across all GFP-Rab expressing cells and it is unlikely that overexpression induces ER movements, consistent with previous reports for other Rabs (Supplementary Figure 4B) (Lu *et al*., 2020; Spits *et al*., 2021). A previous study implicated Rab10 in restructuring the ER in COS7 cells (English and Voeltz, 2013b). We find only a small fraction of Rab6-vesicles colocalized with Rab10 (Supplementary Figure 4C). Furthermore, only ∼50% of Rab6-ER cotransport events had Rab10 present (Supplementary Figure 4D). Taken together, our data suggests that Rab6 is the major post-Golgi Rab responsible for ER hitchhiking.

**Figure 5.**
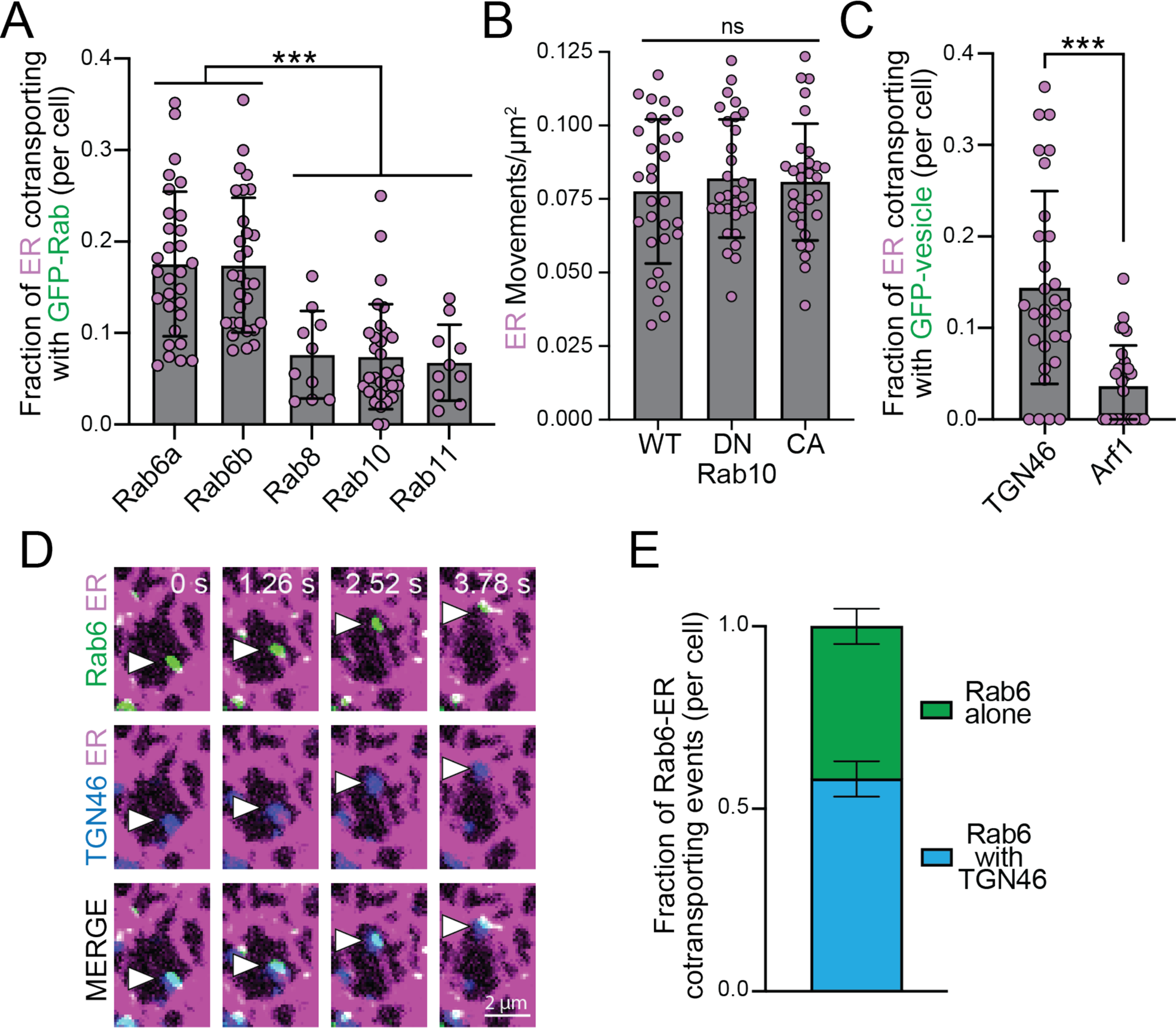
Proximal, but not distal, post-Golgi vesicles participate in ER hitchhiking. (A) ER-Rab cotransport frequency in U2OS cells expressing mCherry-KDEL (ER) and the following GFP-Rab constructs: Rab6a (n=30 cells), Rab6b (n=30), Rab8 (n=10), Rab10 (n=30), and Rab11 (n=10). (Kruskal-wallis with Dunn’s posthoc test, **p<0.01) Data for Rab6a/b is replicated from Figure1B, shown here for comparison. (B) ER movement in U2OS cells expressing mCherry-KDEL (ER) and WT, DN (T23N), and CA (Q68L) GFP-Rab10 constructs (n=30 cells; Brown-Forsythe and Welch ANOVA, ns). (C) Cotransport frequency of TGN46 and Arf1 with ER tubules in U2OS cells expressing mCherry-KDEL (ER) and GFP-TGN46 or mEmerald-Arf1. (n=30 cells per condition; Unpaired t test with Welch’s correction, ***p<0.001). (D) Time-lapse stills of U2OS cells expressing mCherry-KDEL (ER), GFP-TGN46, and Halo-Rab6a (JF646). White arrowheads indicate ER-Rab6a cotransport events with a cotransporting TGN46 molecule. (E) Fraction of Rab6-ER cotransport events with and without cotransporting TGN46 (n=5 technical replicates from 21 analyzed cells). All data represents mean ± SD. ns=not significant.

To further identify the population of Rab6-vesicles participating in ER hitchhiking, we tested cotransport between Rab6 and other post-Golgi proteins. TGN46 marks the trans Golgi network and Golgi-derived vesicles (Luzio *et al*., 1990). We find that ER cotransports with TGN46 to a similar extent as Rab6 (Figure 5C). To determine if this TGN population contains Rab6, we quantified ER cotransport in cells expressing GFP-KDEL (ER), mCherry-TGN46, and Halo-Rab6b. We find that the majority of Rab6-ER cotransporting events were also TGN46-positive (Figure 5D and E). We also tested cotransport between ER tubules and another Golgi-derived vesicle, Arf1, which marks the cis- and trans-Golgi and functions in recruiting coat proteins for vesicle release (Donaldson and Jackson, 2011). We find that Arf1 does not cotransport with ER tubules (Figure 5C). Together, our data indicates that the Rab6-vesicles involved in ER hitchhiking originate from the trans-Golgi network and are a specific subset of Golgi-derived vesicles marked by TGN46.

### ER to Golgi Rab1 participates in ER hitchhiking

Thus far, we have determined that ER hitchhikes on post-Golgi vesicles marked by Rab6. Previous work has shown that Rab1 trafficking between the ER to Golgi and more recently, ER organization during mitosis (Plutner *et al*., 1991; Saraste, 2016; Rollins and Blankenship, 2023). We investigated the participation of Rab1a in ER hitchhiking and find that Rab1a cotransports with ER tubules to a similar extent as Rab6 and Rab7 (Figure 6A and 6B). Furthermore, Rab1a does not colocalize with Rab6 in the periphery of the cell (Supplementary Figure 5A), suggesting this is a new class of vesicles involved in ER hitchhiking. Cells expressing Rab1-DN (S25N) have reduced ER movement compared to WT and GTP-locked (Q70L; constitutively active) expressing cells (Figure 6C and Supplementary Figure 5B). Similar to Rab6, rapalog-induced inhibition of Rab1-vesicle movement significantly reduced ER movement, while rapalog-induced movement towards the cell periphery reorganized the ER to peripheral sites (Figure 6D - 6G). Taken together, our results indicate that ER hitchhikes on Rab1-vesicles and Rab1-vesicle motility is both necessary and sufficient for a subset of ER movements.

**Figure 6.**
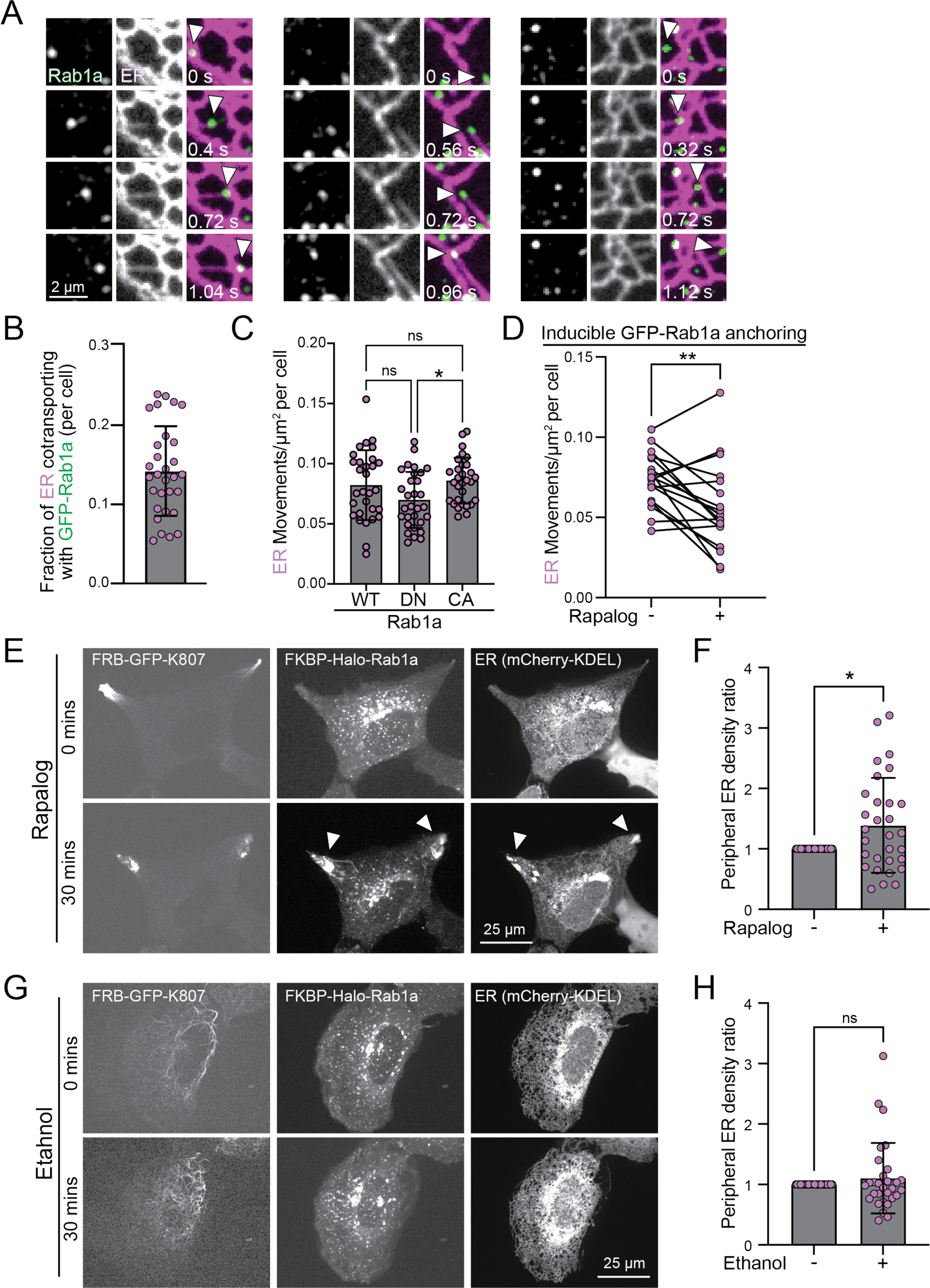
ER to Golgi marker Rab1 participates in ER hitchhiking. (A) Time-lapse stills of U2OS cells expressing GFP-Rab1a and mCherry-KDEL (ER). White arrowheads indicate ER-Rab1 hitchhiking events. (B) ER-Rab1 cotransport frequency in U2OS cells expressing mCherry-KDEL (ER) and GFP-Rab1a. (n=30 cells). (C) ER movement in cells expressing mCherry-KDEL (ER) and either WT, DN (S25N; GDP-locked), or CA (Q70L; GTP-locked) eGFP-Rab1a (n=30 cells; Brown-Forsythe and Welch ANOVA, *p<0.05;). (D) ER movements in U2OS cells before and 30 minutes after treatment with rapalog (n=18 cells; Wilcoxon paired test, *p< 0.0001). (E and G) U2OS cell expressing KIF5B^K807^-Emerald-FRB, 2xFKBP-Halo-Rab1a and mCherry-KDEL (ER) before and after treatment with rapalog (E) or ethanol (G). White arrowheads indicate areas of Rab1-vesicle and ER accumulation. (F and H) Quantification of E and G as a ratio of peripheral to perinuclear ER intensity in the same cell before and after treatment with rapalog (F) (n=30 cells; Wilcoxon paired test, *p<0.05) or ethanol (H) (n=30; Wilcoxon paired test, ns). Data represents mean ± SD.

## DISCUSSION

Proper microtubule-based transport of the endoplasmic reticulum is critical for a variety of functions, including formation of membrane contact sites with other organelles. We investigated an underappreciated mode of ER movement called ‘ER hitchhiking’ which involves peripheral ER tubules tethering to motile Rab-marked vesicles. ER was previously shown to hitchhike on lysosomes Rab5- and Rab7-marked endosomes (Friedman *et al*., 2013; Guo *et al*., 2018; Lu *et al*., 2020; Spits *et al*., 2021; Jang *et al*., 2022). We show here that ER tubules also hitchhike on Golgi-derived Rab6-vesicles independently of ER-Rab7 hitchhiking. Further, these Rab6-vesicles travel on stable MTs, and their motility is both necessary and sufficient to drive a subset of ER movements. Finally, we show that ER hitchhikes on vesicles derived from the TGN and that Rab6 is the only post-Golgi Rab-vesicle that participates in ER hitchhiking. Taken together, these data provide new avenues of research into ER dynamics and raise the question of how prevalent ER hitchhiking is in comparison to other modes of transport. Along these lines, we also find that the ER to Golgi marker Rab1 influences ER movement via hitchhiking.

Endolysosomal vesicles have been shown to reorganize the ER via hitchhiking in response to cellular starvation (Spits *et al*., 2021; Jang *et al*., 2022). This process is regulated by ER membrane protein VAMP-associated protein (VAP) for ER-Rab7 contacts and by MTM1 for ER-Rab5 contact sites (Lu *et al*., 2020; Spits *et al*., 2021; Jang *et al*., 2022). VAP likely serves a dual functional role at ER-endolysosome contact sites. First, VAP aids in lipid transfer between ER and endosome membranes due to its interactions with FFAT-containing lipid transfer proteins (LTPs; Kors *et al*., 2022). Second, VAP acts as a tether at ER and endolysosomal membrane contacts, as knockdown of VAP and its isoforms results in both reduced ER movement and reduced ER-endosome contacts (Di Mattia *et al*., 2018; Spits *et al*., 2021; Kors *et al*., 2022). It is unclear if there is a correlate biological function for ER hitchhiking on Rab6-vesicles, but it is possible that VAP acts as a membrane tether by interacting with FFAT-containing LTPs on the Rab6-vesicle membrane. A major function of ER-Golgi contact sites is non-vesicular lipid transport via LTPs, such as transfer of ceramide from the ER to the Golgi for sphingomyelin synthesis (Hanada, 2010; English and Voeltz, 2013a). Thus, a tethering role for VAP and LTPs on Golgi-derived vesicle membranes is possible.

A function for Rab6-ER hitchhiking may be to coordinate the timing and placement of both compartments where they have overlapping biological roles. For example, ER motility and Rab6 are both critical for focal adhesion growth (Zhang *et al*., 2010; Fourriere *et al*., 2019). Our kinesin-driven rapalog experiment (Figure 4E - G) shows that Rab6 directs ER tubules to specific regions of the cell periphery, reminiscent of directed microtubule-dependent trafficking of Rab6 towards focal adhesions (Fourriere *et al*., 2019). Previous work has also shown that ER tubules constrict mitochondria at fission sites (Friedman *et al*., 2011), and post-Golgi vesicles collaborate with the ER to recruit PI(4P) to ER-mitochondrial contact sites (Nagashima *et al*., 2020). While our data show that mitochondria do not hitchhike on Rab6-vesicles (Figure 1 D - E), it is possible that coordinated movement of Rab6-ER brings the machinery to the correct time and place during mitochondrial division.

Proximal post-Golgi vesicles participating in ER hitchhiking likely originate from the TGN (Figure 5). TGN46 cotransports with ER tubules and colocalizes with Rab6 during hitchhiking events. After exiting the Golgi, TGN46 is trafficked retrogradely from early endosomes and recycling endosomes back to the Golgi (Ghosh *et al*., 1998; Mallet and Maxfield, 1999). Our data is most consistent with Rab6-ER hitchhiking occurring on anterograde Golgi-derived carriers, as demonstrated by (1) the lack of colocalization between Rab6 and Rab7 (Figure 1F and 1G; Supplemental Figure S1D), (2) the absence of Rab11-ER hitchhiking (Figure 5A and Supplemental Figure S4A), and (3) the observation that the majority of ER-Rab6 hitchhiking events move anterogradely (Figure 3E).

We also determined that Rab1-vesicles drive the movement of ER. While a definitive biological function remains elusive, ER-Rab1 hitchhiking may be important during mitosis where ER tubules reorganize around the mitotic spindle in a Rab1-dependent manner (Rollins and Blankenship, 2023). Although our studies were performed in interphase, it is possible that similar mechanisms exist to bridge the Rab1-ER contact site. Another possibility is that ER hitchhiking is important for cargo secretion at ER exit sites (ERES). Secretory cargos are concentrated and exported from the ER to the Golgi in vesicles marked by Rab1 (Westrate *et al*., 2020). The potential function of ER-Rab1 hitchhiking in ERES and ER-to-Golgi transport is an interesting avenue that should be investigated further.

What is the overall prevalence of ER hitchhiking? ER hitchhiking on endolysosomal vesicles contributes to 30-50% of total ER movements (Lu *et al*., 2020; Spits *et al*., 2021). Our data show that ER hitchhiking on Rab1- and Rab6-vesicles are mutually exclusive compartments that each contribute an additional 15-25% of total ER movements. Thus, the majority of ER movements appear to be driven by motile vesicles, as opposed to traditional motor-driven sliding and TAC. This possibility provides an exciting avenue for future research and to understand how cellular demands might dictate which ER movement mechanism and vesicle populations are utilized.

## METHODS

### Cell culture, transfections, and constructs

U2OS cells (ATCC, HTB-96) were maintained on 100 mm tissue-culture treated polystyrene plates (Nest Biotechnology, Cat# 704201), cultured in DMEM (Corning, Cat# MT10013CV) supplemented with 10% fetal bovine serum (FBS; Gibco, Cat# 26140079) and 1% penicillin/streptomycin (Corning, Cat# 30-002-CI), and maintained at 37°C and 5% CO_2_. All transfections were performed using Lipofectamine 2000 (Invitrogen, 11668019) in Opti-MEM (Gibco, Cat# 31985070) according to manufacturer’s instructions. For live-cell imaging experiments, between 50,000 and 100,000 U2OS cells were plated on 35 mm glass-bottom fluorodishes (World Precision Instruments, Cat# FD35-100) or 6-well glass-bottom plates (Cellvis, Cat# P06-1.5H-N) coated with Poly-D-Lysine (PDL) and laminin (PDL: 250 μg/mL, Millipore, Cat# A-003-E; mouse laminin: 4.72 μg/mL, Gibco, Cat# 23017-015). Cells were transiently transfected approximately 24 hours after plating and were imaged ∼24 hours after transfection at 37 ° C and 5% CO_2_. Cells were transfected with 500 ng of each plasmid with the following exceptions: for cotransport between ER, Rab6, and TGN46 (Figure 5D and 5E), U2OS cells were transfected with 250 ng of mCherry-KDEL (ER), 250 ng of Halo-Rab6, and 250 ng mEmerald-TGN46; for ER-Rab6 cotransport on microtubules (Figure 2B and 2C), cells were transfected with 50ng pm-Scarlet-alpha-tubulin, 50 ng mEmerald-Sec61b-C1, and 500 ng pEF-2xFKBP-Halorab6b; for experiments with Halo-tag constructs, transfected U2OS cells were incubated with 200 nM Janelia Fluor 646 HaloTag ligand (Promega, Cat# GA1120) for 30-60 minutes and subsequently washed out with DMEM prior to imaging at 37°C and 5% CO_2_. To visualize mitochondria, 0.2 µl of 1mM Mito-tracker Deep Red was added 60 mins prior to imaging.

mCherry-ER-3 (mCherry-KDEL; Addgene plasmid #55041) and mEmerald-TGNP-N-10 (Addgene plasmid #54279) were gifts from Michael Davidson. mCherry-Sec61-β was a gift from Gia Voeltz (Zurek *et al*., 2011; Addgene plasmid #49155). mEmerald-Sec61β-C1 was a gift from Jennifer Lippincott-Schwartz (Nixon-Abell *et al*., 2016; Addgene plasmid #90992). The following eGFP-Rab plasmids were gifts from Marci Scidmore: eGFP-Rab6a (Rzomp *et al*., 2003; Addgene plasmid #49469), eGFP-Rab6a Q72L (Moorhead *et al*., 2007; Addgene plasmid #49483), eGFP-Rab6a T27N (Moorhead *et al*., 2007; Addgene plasmid #49484), eGFP-Rab6b (Rzomp *et al.,* 2003(Rzomp *et al*., 2003; Addgene plasmid #49470), eGFP-Rab6b Q72L (Addgene plasmid #49889), eGFP-Rab6b T27N (Addgene plasmid # 49890), eGFP-Rab10 (Rzomp *et al*., 2003; Addgene plasmid #49472), eGFP-Rab10 Q68L (Huang *et al*., 2010; Addgene plasmid #49544), eGFP-Rab8 (Huang *et al*., 2010; Addgene plasmid #49543), eGFP-Rab8 T22N (Addgene plasmid #49893), eGFP-Rab13 ((Huang *et al*., 2010; Addgene plasmid #49548), eGFP-Rab14 (Huang *et al*., 2010; Addgene plasmid #49549), and eGFP-Rab14 S25N (Addgene plasmid #49593). eGFP-Rab8 Q67L was a gift from Lei Lu (Madugula and Lu, 2016; Addgene plasmid #86076). GFP-Rab11 WT (Choudhury *et al*., 2002; Addgene plasmid #12674) and GFP-Rab11 DN (Choudhury *et al*., 2002; Addgene plasmid #12674) were gifts from Richard Pagano. eGFP-Rab13 T22N was a gift from Bernardo Mainou (Mainou and Dermody, 2012; Addgene plasmid #110501). eGFP-Rab7a was a gift from Qing Zhong (Sun *et al*., 2010; Addgene plasmid #28047). pmScarlet_alphaTubulin_C1 was a gift from Daphne Bindels (Bindels *et al*., 2017; Addgene plasmid #85045). pSpCas9(BB)-2A-Puro (PX459) V2.0 was a gift from Feng Zhang (Ran *et al*., 2013; Addgne plasmid #62988).

To clone FKBP-Halo-Rab6b and FKBP-Halo-Rab1a, 2x-FKBP-Halotag with either Rab6b cDNA or Rab1a cDNA was subcloned and inserted (using Gibson assembly) into PmeI and EcoRI restriction sites of a pEF vector containing an EF1alpha promoter. p(twist)-KIF5B^K807^-Emerald-FRB with a CMV promoter was synthesized and cloned by Twist Bioscience. For eGFP-Rab10-DN, T23N was cloned into eGFP-Rab10 by site-directed mutagenesis using forward primer 5’ ccggagtggggaagaactgcgtcctttttcg 3’ and reverse primer 5’ cgaaaaaggacgcagttcttccccactccgg 3’.

### Generation of Rab6 double knockouts in U2OS cells using CRISPR/Cas9

To generate the Rab6a/b double knockout via CRISPR, we used a similar approach as previously described (Ran *et al*., 2013). Briefly, 50,000 WT U-2 OS cells were plated on plastic 6-well plates and cultured in DMEM (10% FBS, 5% penicillin/streptomycin) at 37C and 5% CO2. Approximately 24 hours after plating, cells were transiently transfected for one hour with 167ng each of three CRISPR/Cas9 px459 plasmids: targeting Rab6a exon 1 (gRNA primer: 5’ GTCTCCGCCCGTGGACATTG 3’), targeting Rab6b exon 3 (gRNA primer: 5’ GGAAGACGTCTCTGATTACG 3’), and targeting Rab6b exon 5 (gRNA primer: 5’ CTACATCCGGGACTCCACGG 3’). Transfections were performed in OptiMEM with 1.5uL lipofectamine 2000. Approximately 72 hours after transfection, transfected cells were selected by treating with 1ug/mL puromycin for 96 hours total, with media exchanged and fresh puromycin added 48 hours after treatment began. Upon reaching confluency, cells were single cell selected, allowed to grow, and lysed in Laemlli sample buffer with reducing agent. Rab6a/b knockout was confirmed by western blotting using a Rab6 antibody that recognizes both Rab6a and Rab6b isoforms (rabbit anti-Rab6 (ThermoFisher # PA5-22127).

### Microscopy

All microscopy images were acquired using a Nikon Ti-2E inverted microscope with perfect-focus for simultaneous and near-simultaneous (triggered), outfitted with a Yokogawa CSU-X1 spinning disk, two Photometrics Prime BSI sCMOS camera (TwinCam dual camera splitter with filter cubes), and four laser lines (405nm, 488nm, 561nm, and 647nm) with triggered acquisition. The microscope is equipped with a stage-top CO2 and temperature incubator for cell imaging (Oko Labs). The microscope is controlled by a specialized computer that runs Nikon NIS-Elements. Equipped with three objectives: Plan Apo 20x 0.75 NA, 60x 1.4 NA, and 100x 1.45 NA. All imaging experiments in the present study were performed on a 60x oil objective.

All time-lapse images were acquired for a total of 20 sec with no delay. For two-color live-cell imaging, movies were acquired in dual-camera mode at near-simultaneous triggered acquisition (∼50-100ms intervals) except for the mitochondria/Rab6 experiments (Figure 1D and 1E) which was acquired at a 500 ms frame rate. For three-color live-cell imaging experiments, single-camera mode was used with sequential acquisition and no delay; three-color experiments achieved a frame rate of ∼420 msec apart from the assessment of ER-Rab6 cotransport events on microtubules (Figure 2B and 2C), which was ∼900 msec per three-color acquisition.

### Rapalog Heterodimerization Experiments

For all rapalog experiments (Figures 4 and 6), cells were treated with either rapalog A/C heterodimerizer (final concentration 5μM; Takara, Cat# AP21967) or 200 μL ethanol (control). For rapalog-induced Rab-vesicle stoppage experiment (Figure 4A-D and Figure 6D) U2OS cells were transiently transfected with indicated constructs and Janelia Fluor 646 dye was added prior to acquisition. Cells were imaged prior to rapalog or ethanol incubation with 488 nm laser capture (snapshot), 561 nm acquisition for 30 seconds at 250 ms intervals and 640 nm for 20 seconds at 50 msec intervals. The same cells were re-imaged using the same imaging parameters 30-60 mins after incubation with either rapalog or ethanol. For rapalog-induced kinesin-driven Rab-vesicle movement experiment (Figure 4E-I and Figure 6E and 6F), snapshots (taken from the first frame of a 10 s time-lapse video) of all three channels were taken of the same cell before and 30-60 min after incubation with rapalog or ethanol. For the 30 min time-lapse demonstrating re-localization of Rab6-vesicles and ER (Supplemental Movie S3), cells were imaged in single-camera mode with 488nm, 561nm, and 647nm lasers sequentially at 1-min intervals.

To determine efficacy of inducible vesicle anchoring (Figure 4A), FIJI plugin TrackMate was used to track Rab6 vesicle movement (Figure 4B). LoG (Laplacian of Gaussian) detector was used to detect puncta. Object diameter was set to 0.5 microns and quality threshold was adjusted until all puncta were selected. Hyperstack displayer was used as the view window after puncta were detected. Puncta were filtered by maximum intensity to remove puncta that were saturated with signal (frequently marking the Golgi). Simple LAP tracker was used to track puncta movements throughout the movie, with a maximum linking distance of 2.0 microns, a maximum gap-closing distance of 2.0 microns, and a maximum gap-closing fame gap of 1 frame. Tracks were filtered by track displacement > 1 micron to eliminate tracks produced by diffusive vesicle movement. Tracks were colored white uniformly and saved as an overlay on the cell micrograph.

### Data Analysis

All experiments were analyzed using FIJI/ImageJ (NIH). Blind analysis plug-in was used to blind data before analysis. To quantify ER tubule movements, images were bleach corrected and background subtracted, and an ROI of 200-500 μm^2^ of cell area was selected from the cell periphery. Individual ER tubule movements extending from existing ER junctions were counted manually. The total number of ER tubule movements per ROI segment was divided by the area of each ROI to obtain the average number of ER tubule movements per micron^2^ of cell area.

For analyzing the frequency of cotransport events, the Rab channel was processed using Gaussian Blur filter (radius 0.11 µm) followed by Mexican Hat Filter plugin (radius 2.0) Cotransport was measured as the fraction of ER tubules accompanied by a cotransporting puncta (or multiple puncta in the case of three-color experiments) at the ER tip during movement. For microtubule data (Figure 2D and 2E), ER movements on stable microtubules were determined by the presence of microtubules at the onset of cotransport events. The fraction of microtubule-positive co-transporting events and microtubule-absent cotransporting events was given by dividing microtubule-positive or microtubule-absent Rab6 positive-ER tubule movements by total Rab6 positive-ER tubule movements. For analyzing the frequency of mitochondrial cotransport events, average fraction of mitochondrial movements (defined as directed movement >3 frames) accompanied by a cotransporting Rab6 puncta per cell was quantified.

To analyze the speed of Rab6 and the speed and directionality of ER tubules (Figure 2A and Figure 3E), kymographs were generated from time-lapse movies. To generate ER kymographs, the segmented line tool was used to trace (frame by frame) the trajectories of moving individual ER tubules (traced specifically at ER tips), which was subsequently resliced to generate a kymograph (“command” + “/” in ImageJ). For Rab6 puncta, three segmented lines were generated tracing the tracks of Rab6 puncta frame by frame for each contransporting event (before reaching ER tubule, during cotransport with ER tubule, and after dissociation from ER tubule). The speeds of individual particles (for both ER and Rab6) were calculated from the inverse of the slopes of kymograph traces. Only movements >1 second and >0.5 µm were considered. If multiple runs (with pauses in between) were present for a single particle, only the greater of the two run lengths were considered. Directionality (retrograde vs. anterograde) was determined by tubules that clearly moved towards the cell periphery (anterograde) or nucleus (retrograde).

For ER distribution (Figure 4G and 4I and Figure 6F and 6H) FIJI macro CellEdge (Morgan DeSantis, unpublished) was used to calculate peripheral and central ER signal intensity. Briefly, cell outlines were drawn based on GFP-Rab6 or cytosolic GFP signals. Images were background subtracted. Enlarge and band parameters were −4 and 6, respectively. ER distribution was analyzed as a ratio of peripheral ER signal to central ER signal. For enriched vs. non-enriched analysis (Supplementary Figure 3B) ER signal intensity was sorted by enriched and non-enriched Rab6-vesicle signal. Bands of peripheral ER signal generated by Fiji macro CellEdge (Morgan DeSantis, unpublished) were traced with segmented line tool (width =100) to calculate peripheral Rab6-vesicle and ER signal intensity 360 degrees around each cell before and after rapalog addition. Non-enriched Rab6-vesicle intensity was determined from a baseline range calculated from average ± STDEV. Enriched Rab6-vesicle signal intensity was determined by values greater than the maximum baseline range.

### Western Blotting

Cells were trypsinized, spun down at 200 x g, washed with 1x PBS, and lysed in Laemmli sample buffer (Bio-Rad, Cat# 1610737) containing reducing agent (Invitrogen, Cat# NP0009). Samples were incubated at 95°C for 5 minutes. Proteins were separated by SDS-PAGE in 4-12% Bis-Tris Gel (Invitrogen, Cat# NP0321BOX) and transferred onto nitrocellulose (ThermoScientific Cat# 88018) membranesusing Tris-Glycine Transfer buffer with 15% methanol. at 4°C.

### Antibodies

The following primary antibodies were used in this study: Rabbit anti-Rab6a (Invitrogen, Cat# PA5-22127; recognizes both a and b isoforms); rabbit anti-Rab6b (Proteintech, Cat# 10340-I-AP); chicken anti-GFP (Novus, Cat# NB100-161-4-0.02ml); and rabbit anti-GAPDH (Cell Signaling Technology, Cat# 2118S). Secondary antibodies used: anti-Rabbit HRP (Cell Signaling Technology, Cat# 7074S); anti-chicken 488 (Invitrogen, Cat# A11039).

## Acknowledgements

We thank Michael J. Previs for advice on figure illustrations. J.S. is supported by 1R35GM150857 (NIH/NIGMS).

---

**Figure S1.**
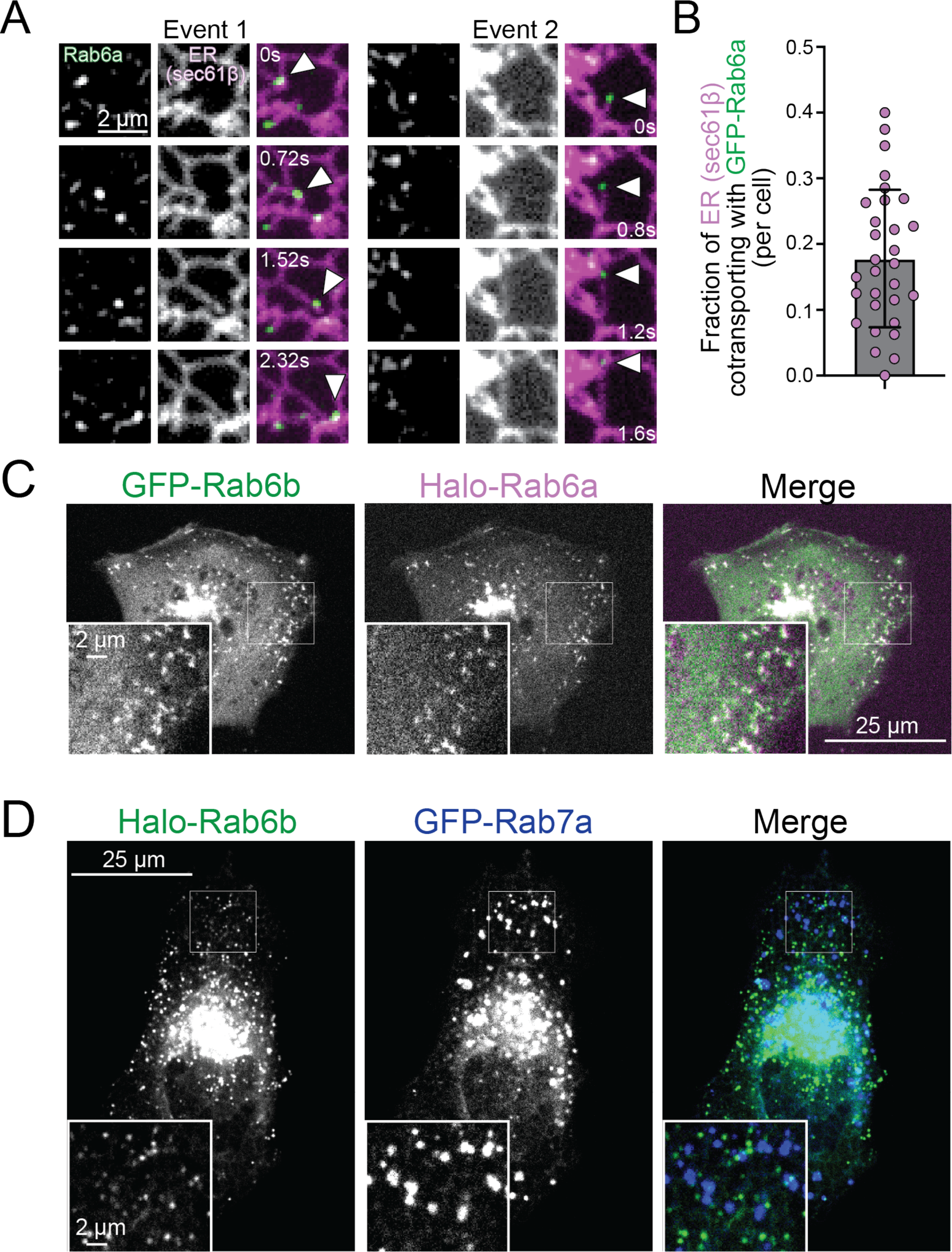
Rab6b colocalizes with Rab6a but not Rab7. (A) Time-lapse stills of U2OS cells expressing mCherry-Sec61β (ER) and eGFP-Rab6a. (B) Cotransport frequency of eGFP-Rab6a with mCherry-Sec61β (ER) (n=30 cells) Data represents mean ± SD. (C) Micrographs of U2OS cells expressing eGFP-Rab6b and Halo-Rab6a(JF646). (D) Micrographs of U2OS cells expressing Halo-Rab6b(JF646) and GFP-Rab7a.

**Figure S2.**
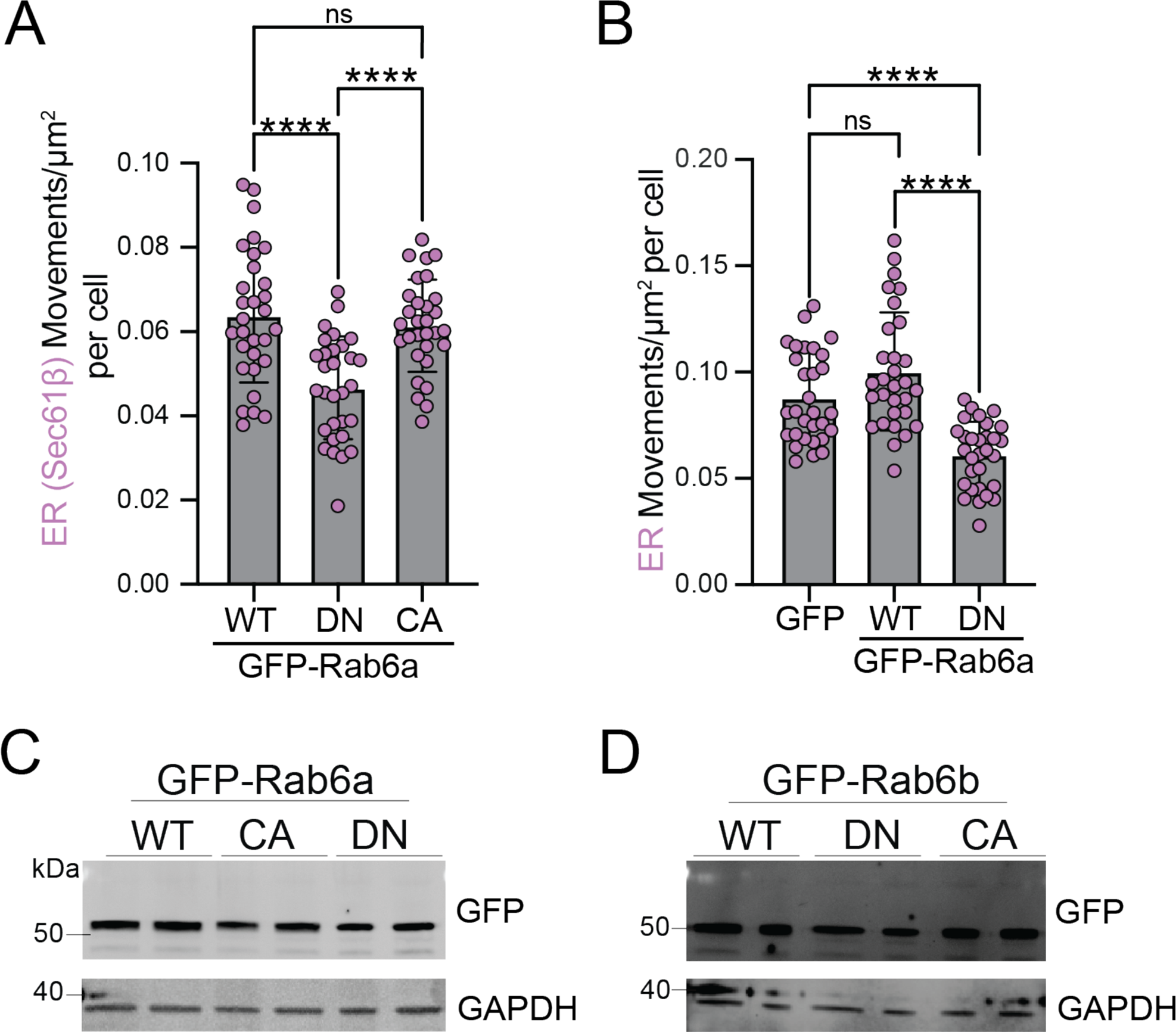
Characterization of Rab6a/b wild-type (WT), dominant-negative (DN), and constitutively-active (CA) constructs. (A and B) U2OS cells expressing WT, DN, and CA eGFP-Rab6a (A) or eGFP-Rab6b (B). (C) ER movement in cells expressing mCherry-Sec61β (ER) and GFP-Rab6a WT, DN, and CA (n=30 cells; Brown-Forsythe and Welch ANOVA, ****p<0.0001). (D) ER movement in cells expressing mCherry-KDEL (ER) and either cytosolic GFP, eGFP-Rab6a WT, or eGFP-Rab6a DN (n=30 cells; Brown-Forsythe and Welch ANOVA, ****p<0.0001. (E and F) Western blots of GFP-Rab6a (E) and GFP-Rab6b (F) expression for WT, DN, and CA constructs in U2OS cells. Data represents mean ± SD.

**Figure S3.**
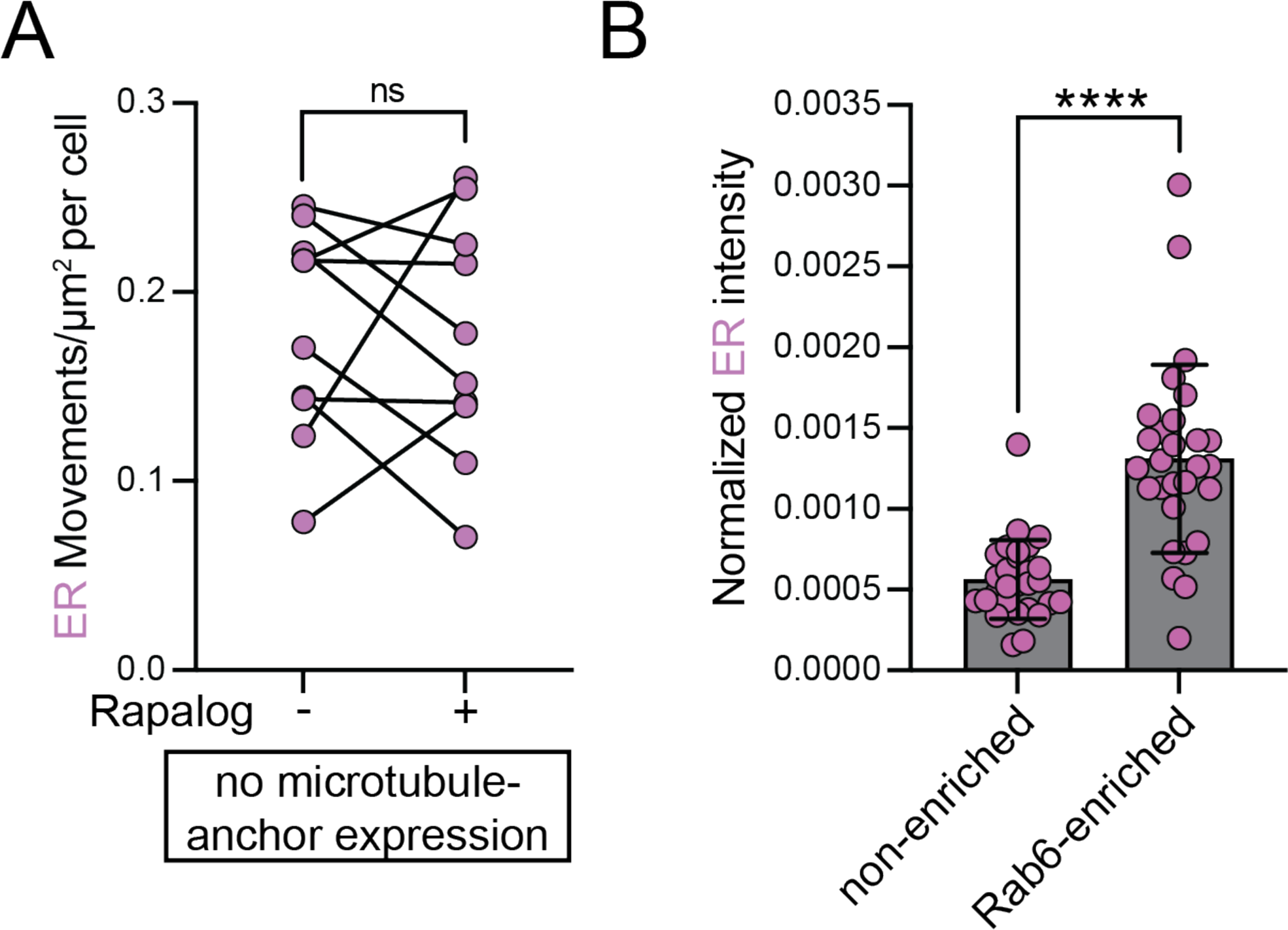
Characterization of rapalog-inducible approach. (A) ER movements in U2OS cells expressing mCherry-KDEL (ER) and 2xFKBP-Halo-Rab6b, lacking FRB-mEmerald-K560^T92N^; (n=10 cells Wilcoxon Paired Test, ns=not significant). (B) ER intensity in U2OS cells expressing KIF5B^K807^-Emerald-FRB, 2xFKBP-Halo-Rab6b and mCherry-KDEL (ER) as ER intensity in peripheral regions with enriched and non-enriched (n=28 cells) Rab6b fluorescence signal (Wilcoxon paired test, ****p<0.0001; Data represents mean ± SD).

**Figure S4.**
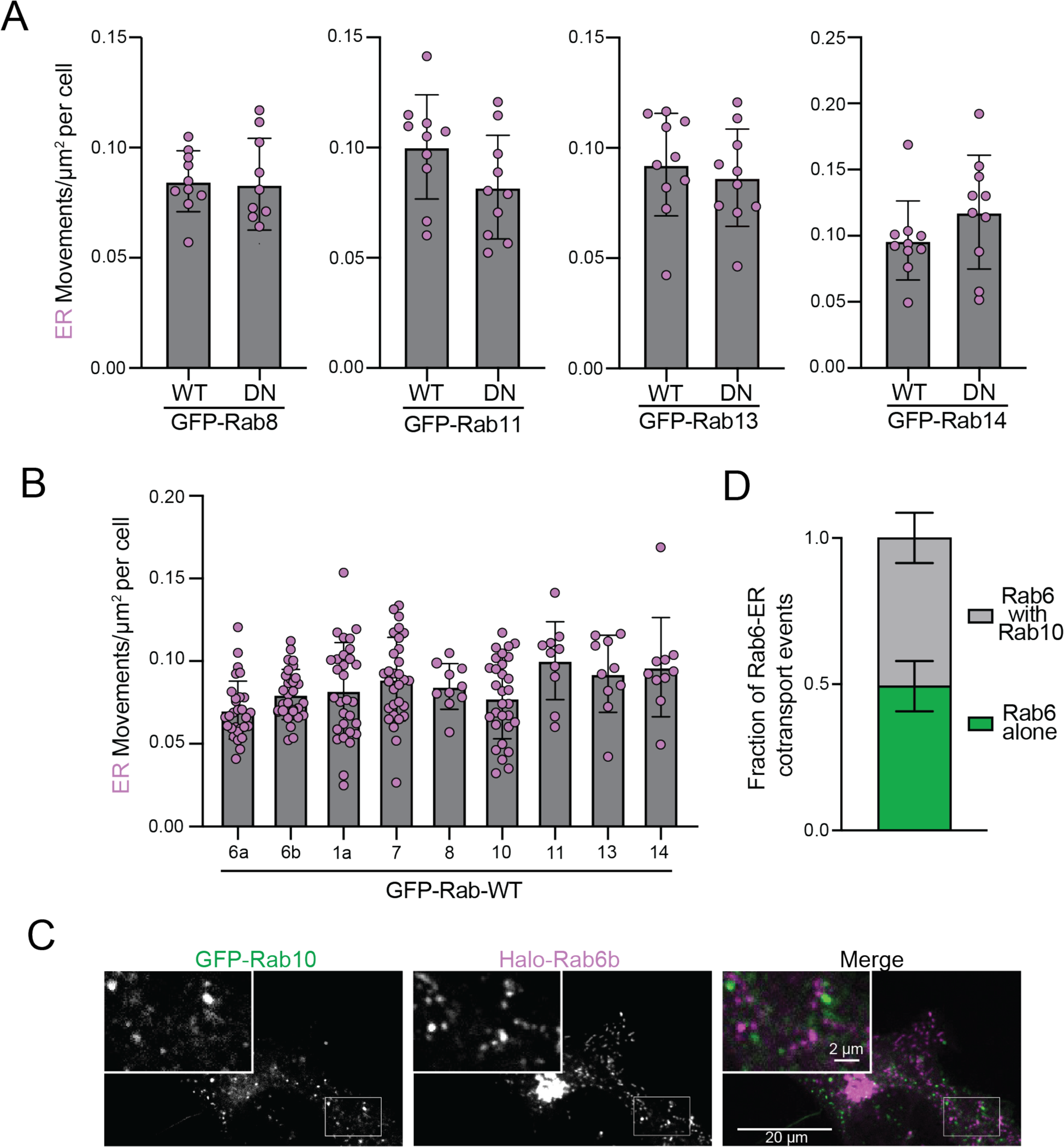
Distal Golgi-derived Rabs do not affect ER movement. (A) ER movement in cells expressing mCherry-KDEL (ER) and either WT or DN constructs of Rab8, Rab11, Rab13, and Rab14 (n=10 cells per condition per Rab; Unpaired t test with Welch’s correction). (B) Total ER movement in cells expressing the following GFP-Rabs: Rab6a (n=30), Rab6b (n=30), Rab1a (n=30), Rab7 (n=30), Rab8 (n=10), Rab10 (n=30), Rab11 (n=10), Rab13 (n=10), and Rab14 (n=10); all n refers to number of cells (C) U2OS cell expressing GFP-Rab10 and Halo-Rab6b (JF646). (D) Fraction of ER-Rab6b cotransport events with GFP-Rab10 (N=5 technical replicates; Data represents mean ± SEM).

**Figure S5.**
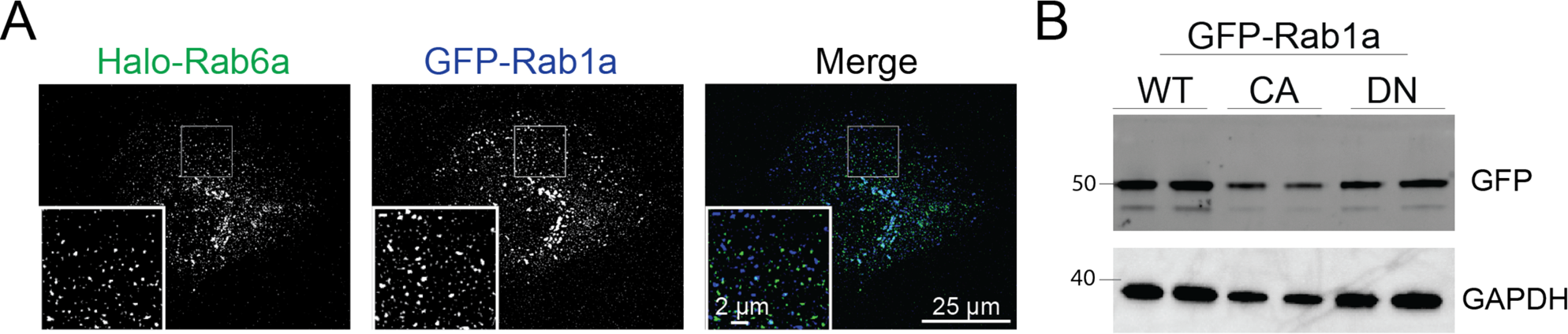
GFP-Rab1 expression and colocalization with Rab6 (A) U2OS cells coexpressing Halo-Rab6a(JF646) and GFP-Rab1a. (B) Western blot of GFP-Rab1a expression in U2OS cells for WT, DN, and CA constructs.

## MOVIE LEGENDS

**Movie S1.** ER cotransports with Rab6-marked vesicles. Merged movie of GFP-Rab6a (green) and mCherry-KDEL (magenta). Green arrows indicate Rab6-vesicles pre-, during- and post-cotransporting. Magenta arrows indicate onset of ER hitchhiking event. Images were acquired on an inverted spinning-disk microscope (Nikon) using dual camera, two-color near-simultaneous(triggered) acquisition.

**Movie S2.** ER cotransports with Rab6-marked vesicles in the absence of Rab7-marked late endosomes. Merged movie of JF646-dyed Halo-Rab6 (yellow), GFP-Rab7 (blue) and mCherry-KDEL (white). Yellow arrows indicate Rab6-ER cotransporting event. Images were acquired on an inverted spinning-disk microscope (Nikon) using single-camera sequential acquisition.

**Movie S3.** Kinesin-driven Rab6-vesicles towards the cell periphery rearranges the ER. Merged movie of JF646-dyed 2xFKBP-Halo-Rab6b (green) and mCherry-KDEL (magenta) in cells expressing KIF5B^K807^-mEmerald-FRB. White circle indicates addition of rapalog. Images were acquired on an inverted spinning-disk microscope (Nikon).

